# Is the 25-year hepatitis C marathon coming to an end to declare victory?

**DOI:** 10.1101/115378

**Authors:** Khulood Ahmed, Ashraf Almashhrawi, Jamal A. Ibdah, Veysel Tahan

## Abstract

Hepatitis C virus (HCV) which was originally recognized as posttransfusion non-A, non-B hepatitis has been a major global health problem affecting 3% of the world population. Interferon/peginterferon and ribavirin combination therapy was the backbone of chronic HCV therapy for two decades of the journey. However, the interferon based treatment success rate was around 50% with many side effects. Many chronic HCV patients with psychiatric diseases, or even cytopenias, were ineligible for HCV treatment. Now, we no longer need any injectable medicine. New direct-acting antiviral agents against HCV allowed the advance of interferon-free and ribavirin-free oral regimens with high rates of response and tolerability. The cost of the medications should not be a barrier to their access in certain parts of the world. While we are getting closer, we should still focus on preventing the spread of the disease, screening and delivering the cure globally to those in need. In the near future, development of an effective vaccine against HCV would make it possible to eradicate HCV infection worldwide completely.

## INTRODUCTION

It has been less than three decades since the discovery of hepatitis C virus (HCV) in 1989 by Choo and colleagues (1). The process of discovering the virus was very daunting as described in Dr. Houghton’s paper (2). Recognized as the reason behind non-A non-B hepatitis, the big picture of the health and financial burden this virus would have caused became clear. With the high sustained virologic responses reported recently with the use of direct acting, soon will be forgotten the miserable quality of life patients of hepatitis C have had to endure with the not as effective and with unpleasant side effects interferon-based treatments. Not until five years ago when direct acting agents, protease inhibitors telaprevir (Incivec, Vertex) and boceprevir (Victrelis, Merck) were approved by the FDA, had we started seeing sustained virologic response (SVR) rates above 70%. Since then, many direct acting agents have been approved with SVR rates above 90%. While very promising, challenges for treatment, such as access to medications and healthcare management, remain widely spread.

## NATURAL HISTORY OF HEPATITIS C

After the acute infection, only 15%-25% of the patients get cured spontaneously and 75%-85% develop chronic infection with the diagnosis made if viremia persists 6 months from the onset. Chronic inflammation will lead eventually to structural damage or fibrosis which, as it progresses, will lead to cirrhosis. From those chronically infected, 10% to 15% develop cirrhosis (3). Progression of fibrosis can be influenced by host factors (such as older age at time of infection, male gender, coinfection with human immunodeficiency virus (HIV) or hepatitis B virus (HBV), immunosuppression, insulin resistance, non-alcoholic steatohepatitis, hemochromatosis, schistosomiasis, and the grade and stage on the liver biopsy) as well as external factors (such as excessive alcohol drinking).

Liver biopsy remains the gold standard for the grading and staging of chronic hepatitis C. The grade, which reflects the inflammation activity, is determined by the severity of mononuclear inflammatory cells around the portal areas and by necrosis of the hepatocytes. The stage reflects the extent of the fibrosis which ranges from absent to mild or advanced in case of bridging fibrosis (fibrosis extending from a portal tract to another) or cirrhosis (fibrosis closing up in circles forming nodules).

Deaths usually are caused by complications of cirrhosis such as ascites, variceal bleeding, hepatic encephalopathy, hepatorenal syndrome, and hepatocellular carcinoma (HCC). Unfortunately, the disease can progress silently until it is advanced and complications ensue. For compensated cirrhosis, 3, 5, and 10-year survival rates were 96%, 91%, and 79%, respectively (4). The 5-year survival rate drops to 50% once decompensated (4). HCC risk increased 17-fold in HCV-infected patients (5). This risk appears to have decreased in those with a sustained viral response rather than non-responders to interferon treatment (6). The rates of progression to cirrhosis and HCC have been variable with a mean time to cirrhosis estimated at 20 years (3, 7). HCC can develop at a rate of 1% to 4% per year (8-11).

## EPIDEMIOLOGY OF HEPATITIS C

The prevalence of hepatitis C virus antibody in the USA is 1.6% (12). In 1999–2002, the highest prevalence (4.3%) was in people 40 to 49 years of age (12) and two-thirds of those infected were born between 1945- 1965. In 2007, HCV infection was associated with an estimated 15,000 deaths in the USA. (13). This number has risen to 19,695 in 2014 and this is now thought of as only a fraction of the actual number (14). Decompensated chronic HCV is the most common indication for liver transplantation in the USA (15, 16). Chronic hepatitis C is a leading cause of hepatocellular carcinoma (17). The incidence of acute hepatitis C in the USA was estimated to be 180,000 cases per year in the mid-1980s, but declined to approximately 30,000 new cases per year in 1995 (18), and to 16,000 cases in 2009 (19). In a more recent surveillance, the incidence of acute hepatitis C in the USA has been on the rise since 2011. The estimated number on the actual new cases was 16,500 in 2011 and has risen to 30,500 in 2014 (14).

## SCREENING FOR HEPATITIS C

Because of the lack of symptoms in compensated disease and because 75% of patients chronically infected with hepatitis C are unaware of their infection (20), screening can help identify those infected before their disease progresses to a late stage. The U.S. Preventive Services Task Force (USPSTF) has found evidence that screening high risk population and one time screening of those born between 1945 and 1965 is of moderate benefit. The risk of stigmatization appeared small and, although there was evidence for harm from liver biopsy of 1% bleeding risk and < 0.2 *%* death risk from liver biopsy, the use of liver biopsy to guide the management is becoming less. In light of the availability of effective antiviral agents that have rare and self-limited side effects, identifying patients with chronic hepatitis C and treating them is probably of benefit (21).

There is adequate evidence that anti–HCV antibody testing followed by confirmatory polymerase chain reaction testing accurately detects chronic HCV infection. The number needed to screen to identify 1 case of chronic hepatitis c in a high risk population, such as past or present injection drug use, sex with an injection drug user, or blood transfusion before 1992, is < 20 persons and anti–HCV antibody testing is associated with high sensitivity (>90%) (22). There is also evidence that different noninvasive tests have good diagnostic accuracy in detecting fibrosis (23).

## IS THERE A VACCINE YET?

Unlike hepatitis A and B, no vaccine is available to protect against hepatitis C. The high variability among different strains and the fast rate at which mutations can develop made it very challenging to create an effective vaccine (24). Several attempts are currently made to create a vaccine either by directing efforts at a relatively stable glycoprotein that is used by the virus to invade liver cells (24).

## ASSAYS

Testing for hepatitis C has improved over the years. Anti-hepatitis C tests are accurate and with high sensitivity >90% in high risk groups. The CDC recommends one of three immunoassays; two enzyme immunoassays (EIA) (Abbott HCV EIA 2.0, Abbott Laboratories, Abbott Park, Illinois, and Ortho^®^ HCV Version 3.0 ELISA, Ortho-Clinical Diagnostics, Raritan, New Jersey) and one enhanced chemiluminescence immunoassay (CIA) (Vitros^®^ Anti-HCV assay, Ortho-Clinical Diagnostics, Raritan, New Jersey) for the initial screening (25). The OraQuick test has been also approved by the FDA for the initial screening in 2011 (26). A reactive initial test should be followed by a confirmatory nucleic acid test (NAT) where the plasma is tested to detect (qualitative) or detect and quantify (quantitative) hepatitis C RNA. If HCV RNA is detected, that indicates active hepatitis C infection. If HCV RNA is not detected, that indicates a false positive HCV antibody test or resolved infection (27). Table 1 shows FDA approved anti-HCV tests (28).

**Table 1:** FDA approved Anti-HCV tests^28^

| **Table 1: FDA approved Anti-HCV tests** <sup>28</sup> |  |  |
| --- | --- | --- |
| **Abbott HCV EIA 2.0** | Abbott Laboratories, AbbottPark, IL | EIA (Manual) |
| **Advia Centaur HCV** | Siemens, Malvern, PA, USA | CIA (Automated) |
| **ARCHITECT Anti-HCV** | Abbott Laboratories, AbbottPark, IL | CMIA (Automated) |
| **AxSYM Anti-HCV** | Abbott Laboratories, AbbottPark, IL | MEIA (Automated) |
| **OraQuick Rapid Test** | OraSure Technologies, Bethlehem, PA | Immunochromatographic (Manual) |
| **Ortho HCV Version 3.0 EIA** | Ortho | EIA (Manual) |
| **VITROS Anti-HCV** | Ortho | CIA (Automated) |
| Anti-HCV = HCV antibody; EIA = enzyme immunoassay; CIA = chemiluminescent immunoassay; MEIA= microparticle enzyme immunoassay; CMIA = chemiluminescent microparticle immunoassay |  |  |

## MILESTONES IN THE HISTORY OF HEPATITIS C

1975 Non-A, non-B hepatitis was first described (29, 30).

1989 Randomized controlled trials were carried out using interferon alpha to treat non-A, non-B hepatitis (31-33).

1989 Hepatitis C virus was identified (1).

1991 Ribavirin is used as a monotherapy for chronic hepatitis C (34, 35).

1995 The combination of interferon alpha and ribavirin were tested (36, 37).

1996 Hepatitis C serine protease structure was published (38).

1998 First randomized double-blind, placebo controlled study using recombinant interferon alpha alone or in combination with ribavirin (39, 40).

1999 Structure of hepatitis C RNA- dependent RNA polymerase NS5B was identified (41, 42).

2001 Pegylated interferon alpha and ribavirin were used in trials (43, 44).

2005 Structure of NS5A was published (45).

2011 First direct acting agents: Protease inhibitors were used in combination with pegylated interferon and ribavirin to treat hepatitis C genotype 1 (46, 47).

2012 Pilot studies using combinations of direct-acting antiviral drugs without interferon (48).

2014- Several direct acting antiviral medications were released to the market to treat different hepatitis C genotypes with sustained virologic response exceeding 90% and with better tolerability.

## DIRECT ACTING AGENTS ERA

In 2011, telaprevir and boceprevir became available as the first direct acting antivirals for the treatment of chronic hepatitis C with variable sustained virologic responses (SVRs) for the different genotypes. While the use of the protease inhibitors was a great milestone in the journey of treating hepatitis C with SVRs above 70%, it had limiting factors, such as the need to use in combination with pegylated interferon and ribavirin, and their limitations in treating those with decompensated cirrhosis. pegylated interferon and ribavirin have exerted their antiviral activities by modulating the host immunity rather than directly inhibiting the virus. With the better understanding of the hepatitis C virus non-structural proteins, agents that inhibited those proteins have shown promise by directly disabling the life cycle of the virus.

To overcome those species with resistance against the various agents, combination therapy with agents attacking different vital functions became available. Several trials have been conducted with many drugs and combinations of drugs tested in patients with different genotypes and subtypes as well as groups who were treatment naive and those with experience. Defining cure by achieving SVR, defined as the absence of virus detection with an acceptable HCV RNA assay at 12 weeks after the end of the treatment, therapy outcomes were compared in those with and without cirrhosis, and between those with and without disease decompensation. While some data is still lacking in different special groups, we are learning more about how to treat those with comorbidities such as coinfection with HIV or those with renal disease. Most of the regimens available at this time are interferon –free regimens and all oral medications with very limited side effects and with SVRs > 85% and in many cases > 95%. Table 2 summarizes the latest recommendations for treating the different HCV genotypes in different groups of patients (49-58).

**Table 2:** AASLD/IDSA Guideline Recommendations: Genotypes 1,2,3,4,5 and 6 HCV^58^

| HCV Genotype | Cirrhosis | Prior Tx | Recommended Regimen | Alternative Regimen | Notes |
| --- | --- | --- | --- | --- | --- |
| 1a | No |  | LDV/SOF 12 wks |  |  |
|  |  |  | DCV+ SOF 12 wks |  |  |
|  |  |  | SMV+SOF 12 wks |  |  |
|  |  |  | SOF/VEL 12 wks |  |  |
|  |  |  | GZR/EBR 12 wks |  | NS5A RAVs absent |
|  |  |  |  | GZR/EBR 16 wks + RBV | NS5A RAVs present |
|  |  |  | OBV/PTV/RTV+DSV 12 wks +RBV |  |  |
| 1b | No |  | LDV/SOF 12 wks |  |  |
|  |  |  | DCV+ SOF 12 wks |  |  |
|  |  |  | SMV+SOF 12 wks |  |  |
|  |  |  | SOF/VEL 12 wks |  |  |
|  |  |  | GZR/EBR 12 wks |  |  |
|  |  |  | OBV/PTV/RTV+ DSV 12 wks |  |  |
| 1a | Compensated | Naive | LDV/SOF 12 wks |  |  |
|  |  |  |  | DCV+ SOF 24 wks +/- RBV |  |
|  |  |  |  | SMV+SOF 24 wks +/- RBV | No Q80K |
|  |  |  | SOF/VEL 12 wks |  |  |
|  |  |  | GZR/EBR 12 wks |  | NS5A RAVs absent |
|  |  |  |  | GZR/EBR 16 wks + RBV | NS5A RAVs present |
|  |  |  |  | OBV/PTV/RTV+DSV 24 wks +RBV |  |
| 1a | Compensated | PR exp | LDV/SOF 12 wks +RBV or 24 wks |  |  |
|  |  |  |  | DCV+ SOF 24 wks +/- RBV |  |
|  |  |  |  | SMV+SOF 24 wks +/- RBV | No Q80K |
|  |  |  | SOF/VEL 12 wks |  |  |
|  |  |  | GZR/EBR 12 wks |  | NS5A RAVs absent |
|  |  |  |  | GZR/EBR 16 wks + RBV | NS5A RAVs present |
|  |  |  |  | OBV/PTV/RTV+DSV 24 wks +RBV |  |
| 1b | Compensated | Naive | LDV/SOF 12 wks |  |  |
|  |  |  |  | DCV+ SOF 24 wks +/- RBV |  |
|  |  |  |  | SMV+SOF 24 wks +/- RBV |  |
|  |  |  | SOF/VEL 12 wks |  |  |
|  |  |  | GZR/EBR 12 wks |  |  |
|  |  |  | OBV/PTV/RTV+DSV 12 wks |  |  |
| 1b | Compensated | PR exp | LDV/SOF 12 wks +RBV or 24 wks |  |  |
|  |  |  |  | DCV+ SOF 24 wks +/- RBV |  |
|  |  |  |  | SMV+SOF 24 wks +/- RBV |  |
|  |  |  | SOF/VEL 12 wks |  |  |
|  |  |  | GZR/EBR 12 wks |  |  |
|  |  |  | OBV/PTV/RTV+DSV 12 wks |  |  |
| 1a or 1b | Decompensated | Naive or exp | SOF/VEL 12 wks + RBV |  | Child-Pugh B or C |
| 2 | No | Naive | SOF/VEL 12 wks |  |  |
|  |  |  |  | SOF + DCV 12 wks |  |
| 2 | No | PR exp | SOF/VEL 12 wks |  |  |
|  |  |  |  | SOF + DCV 12 wks |  |
| 2 | No | SR exp | DCV+ SOF 24 wks +/- RBV |  |  |
|  |  |  | SOF/VEL 12 wks + RBV |  |  |
| 2 | Compensated | Naive | SOF/VEL 12 wks |  |  |
|  |  |  |  | SOF + DCV 16-24 wks |  |
| 2 | Compensated | PR exp | SOF/VEL 12 wks |  |  |
|  |  |  |  | SOF + DCV 16-24 wks |  |
| 2 | Compensated | SR exp | DCV+ SOF 24 wks +/- RBV |  |  |
|  |  |  | SOF/VEL 12 wks + RBV |  |  |
| 2 | Decompensated | Naive or exp | SOF/VEL 12 wks + WB RBV |  | Child-Pugh B or C |
|  |  |  | DCV+ SOF 12 wks + low initial dose RBV |  | Child-Pugh B or C |
| 3 | No | Naive | SOF + DCV 12 wks |  |  |
|  |  |  | SOF/VEL 12 wks |  |  |
| 3 | No | PR exp | SOF + DCV 12 wks |  |  |
|  |  |  | SOF/VEL 12 wks |  |  |
| 3 | No | SR exp | DCV+ SOF 24 wks + RBV |  |  |
|  |  |  | SOF/VEL 12 wks + RBV |  |  |
| 3 | Compensated | Naive | SOF/VEL 12 wks |  |  |
|  |  |  | SOF + DCV 24 wks +/- RBV |  |  |
| 3 | Compensated | PR exp | SOF/VEL 12 wks + RBV |  |  |
|  |  |  | SOF + DCV 24 wks + RBV |  |  |
| 3 | Compensated | SR exp | SOF + DCV 24 wks + RBV |  |  |
|  |  |  | SOF/VEL 12 wks + RBV |  |  |
| 3 | Decompensated | Naive or exp | SOF/VEL 12 wks + WB RBV |  | Child-Pugh B or C |
|  |  |  | DCV+ SOF 12 wks + low initial dose RBV |  | Child-Pugh B or C |
| 4 | No cirrhosis or compensated | Naive | SOF/LDV 12 wks |  |  |
|  |  |  | OBV/PTV/RTV 12 wks + RBV |  |  |
|  |  |  | GRZ/EBV 12 wks |  |  |
|  |  |  | SOF/VEL 12 wks |  |  |
| 4 | No | PR exp | SOF/LDV 12 wks |  |  |
|  |  |  | OBV/PTV/RTV 12 wks + RBV |  |  |
|  |  |  | GRZ/EBV 12 wks |  | Relapse |
|  |  |  | GRZ/EBV 16 wks + RBV |  | Others |
|  |  |  | SOF/VEL 12 wks |  |  |
| 4 | Compensated | PR exp | SOF/LDV 12 wks + RBV | SOF/LDV 24 wks |  |
|  |  |  | OBV/PTV/RTV 12 wks + RBV |  |  |
|  |  |  | GRZ/EBV 12 wks |  | Relapse |
|  |  |  | GRZ/EBV 16 wks + RBV |  | Others |
|  |  |  | SOF/VEL 12 wks |  |  |
| 5 or 6 | No cirrhosis or compensated | Naive or PR exp | SOF/LDV 12 wks |  |  |
|  |  |  | SOF/VEL 12 wks |  |  |

AASLD, American Association for the Study of Liver Diseases; DCV, daclatasvir; DSV, dasabuvir; EBR, elbasvir; GT, genotype; GZR, grazoprevir; LDV, ledipasvir; OBV, ombitasvir; PR, peginterferon/ribavirin; PTV, paritaprevir; RAV, resistance associated variant; RBV, ribavirin; RTV, ritonavir; SMV, simeprevir; SOF, sofosbuvir; VEL, velpatasvir; WB, weight-based; SR, sofosbuvir/ribavirin; exp, experienced; Tx, treatment; wks, weeks.

## SO DID WE FIND THE IDEAL CURE?

With manufacturing highly potent medications, we may be winning a battle against hepatitis C, but are we winning the war? The challenge in eradicating the disease lies in delivering and administering those medications to the appropriate patients and ensure their compliance with the treatment and follow up. Patient access to the medications is limited by cost, lack of healthcare services, and ignorance. While the AASLD recommends treating everybody with chronic hepatitis C, third party payers may restrict treatment to those with advanced disease. Payers may also limit the choice of medications authorized. Many healthcare providers choose not to treat hepatitis C because of the perception that such treatment is complex or at least time and resources consuming. Some providers and insurers will decline those with ongoing illicit drug or alcohol use. Many patients, an estimated 75% of those infected, do not know they have the disease and, therefore, do not seek the appropriate care. While such hindrances have been there all along, the challenge seems to be shifting more towards socioeconomic nature.

## WHAT IS NEXT?

As more direct acting agents are coming out of the pipeline, healthcare managers will have to face the major task of making those medicines available to chronic hepatitis C patients. Spreading awareness about the need for screening and for treatment if infected not only in the population at large, but also among the healthcare providers, will help identifying more cases of infection and help provide those with the appropriate treatment. One of the efforts that is successfully dismantling some of those barriers in the Extended Community Healthcare Outcomes project (ECHO). Launched in New Mexico in 1993, the project has grown to encompass several national and international hubs. These hubs provide healthcare providers with the appropriate educational and coaching resources to empower them with the knowledge and with the How-To guidance to treat patients in their communities where no specialized care is available. This project has helped many patients in rural areas to receive treatment without the need to travel out of their own towns (59, 60). The project provides healthcare providers with direct access to specialized knowledge and provides them with step-by-step coaching which helps in alleviating the misperceptions about the complexity of the treatment and, hence, recruiting more providers in the war against hepatitis C.

On the preventative front, efforts should be also directed towards studying the increasing incidence of hepatitis C acute infection to identify the newer trends behind this surge and to try to eliminate them. While treating acute hepatitis C infection is still not recommended, we may need to revisit the guidelines to facilitate earlier treatment, particularly now that we have highly potent medications with few tolerable side effects.

Finally, efforts toward developing effective vaccines should be boosted as history tells us that most of success stories in eradicating infectious illness were made possible largely because of vaccines against the offending pathogen.

## CONCLUSION

While we are getting closer, it is still early to declare victory against hepatitis C. We are armed with better ammunition, but we need to do better job in developing strategies that not only deliver cure to those in need, but also prevents the spread of the disease by educating the population about the risk factors for contracting the disease and how to avoid them, identifying the undiagnosed, and providing early treatment to those in need.

